# Multimodal precision MRI of the individual human brain at ultra-high fields

**DOI:** 10.1101/2024.06.17.596303

**Authors:** Donna Gift Cabalo, Ilana Ruth Leppert, Risa Thevakumaran, Jordan DeKraker, Youngeun Hwang, Jessica Royer, Valeria Kebets, Shahin Tavakol, Yezhou Wang, Yigu Zhou, Oualid Benkarim, Nicole Eichert, Casey Paquola, Julien Doyon, Christine Lucas Tardif, David Rudko, Jonathan Smallwood, Raul Rodriguez-Cruces, Boris C. Bernhardt

## Abstract

Multimodal neuroimaging, in particular magnetic resonance imaging (MRI), allows for non-invasive examination of human brain structure and function across multiple scales. Precision neuroimaging builds upon this foundation, enabling the mapping of brain structure, function, and connectivity patterns with high fidelity in single individuals. Highfield MRI, operating at magnetic field strengths of 7 Tesla (T) or higher, increases signal-to-noise ratio and opens up possibilities for gains spatial resolution. Here, we share a multimodal Precision Neuroimaging and Connectomics (PNI) 7T MRI dataset. Ten healthy individuals underwent a comprehensive MRI protocol, including T1 relaxometry, magnetization transfer imaging, T2*-weighted imaging, diffusion MRI, and multi-state functional MRI paradigms, aggregated across three imaging sessions. Alongside anonymized raw MRI data, we release cortex-wide connectomes from different modalities across multiple parcellation scales, and supply “gradients” that compactly characterize spatial patterning of cortical organization. Our precision MRI dataset will advance our understanding of structure-function relationships in the individual human brain and is publicly available via the Open Science Framework.

## Background and Summary

Neuroimaging has advanced our understanding of the human brain by allowing non-invasive and large-scale examination of structural and functional brain networks. ^1,2^ Nevertheless, most human MRI research collect limited individual-specific data in brief scanning sessions.^3,4^ Consequently, standard neuroimaging studies predominantly centered around group-averaged data that, while revealing fundamental principles of brain organization, limit the specificity and clinical utility of MRI.^4–6^ Precision neuroimaging, which prioritizes individualized mapping of brain structure and function through the use of repeated and prolonged scans,^5,7^ has emerged as a powerful approach to address this issue. By scanning each individual in long and often repeated sessions, precision neuroimaging provides sufficient signal and data quality to study individuals in their own “native” space.^5,7^ This personalized approach ensures reliable estimates and captures fine-grained organization of networks without the additional blurring of inter-individual variability.^7^ Moreover, averaging structural sequences across multiple scans enhances signal-to-noise ratio (SNR), facilitating the integration of structure and function to interrogate their relationship.

Multimodal neuroimaging approaches hold promise in advancing the understanding of both healthy and diseased states by providing a comprehensive view of individual brains,^8^ thus improving specificity in MRI phenotyping. Multimodal structural imaging often capitalizes on diffusion MRI tractography to examine large-scale connectome architecture and are often complemented with measures of cortical thickness or geodesic distance.^9^ The ability of structural MRI to interrogate brain tissue can be augmented by using quantitative MRI sequences, enriching the biophysical characterization of inter-regional heterogeneity and inter-individual differences in the human brain. ^10^ ^11,12^ ^13^ Notably, T1 relaxation mapping distinguishes highly myelinated regions from less myelinated ones,^10,13^ hence facilitating the *in vivo* investigation of intracortical microstructure and its cognitive implications.^11,12^ Echoing *post-mortem* neuroanatomical studies, this method has revealed smooth transitions in cortical laminar architecture, from sensory and motor cortices to paralimbic circuits.^12,14^ To characterize functional architectures, employing multi-state functional MRI (fMRI) allows investigation of how different brain networks interact and reconfigure under varying circumstances.^15^ For instance, dense temporal sampling of resting-state functional MRI data allows for detailed and reliable characterization of intrinsic functional networks and can help better understand the idiosyncrasy of heteromodal systems.^6,16^

Complementing task-free investigations with task-based fMRI can provide insights into brain responses to specific stimuli or cognitive tasks, providing detailed information on functional specialization.^17^ In this context, the use of movie watching paradigms^18^ have emerged as a valuable tool, allowing examination of synchronization of low-level brain activity and facilitating the identification of individual differences. Movies, closely resembling real life experiences, provide an ecologically valid alternative to both rs-fMRI, which may lack constraints, and task-fMRI, which emphasizes the activity of unique neural circuits.^19^ Additionally, movies help mitigate participant head motion while improving arousal and compliance.^18,20^

Harnessing highfield neuroimaging at magnetic field strengths of 7 Tesla or above can further enhance spatial resolution and sensitivity to blood oxygenation level dependent (BOLD) contrast.^21–23^ Moreover, multi-echo fMRI, a technique that addresses the indeterminacy problem of signal sources, offers improved signal fidelity and interpretability compared to single-echo fMRI.^24^ By acquiring multiple echo images per slice and modeling T2* decay at every voxel, multi-echo fMRI distinguishes brain activity from artifactual constituents.^24,25^ Multiband acceleration enables simultaneous acquisition of planar imaging slices, reducing imaging times.^25,26^ The increased specificity provided by multimodal, highfield, precision MRI therefore enables a targeted delineation of cortical network organization,^27^ encompassing microstructure, connectivity, and function.^26,28^

Recent advancements in neuroimaging and network neuroscience have facilitated the study of large-scale spatial trends in brain structure and function, commonly known as *gradients*.^29–33^ These gradients span various aspects of brain organization including structural^34–36^ and functional connectivity^31,37–39^, task-based investigations^17,40^ cortical morphology and microstructure,^11,12,41,42^ indicating converging spatial trends.^29,32^ For example, analyses of intrinsic functional connectivity gradients have identified a principal gradient distinguishing sensorimotor systems from transmodal networks,^31^ consistent with established cortical hierarchy models.^43^ This gradient’s pattern also reflects geodesic distance measures between sensory and transmodal regions, suggesting a mechanism enabling transmodal networks to support higher cognitive functions independent of immediate sensory input.^44,45^ Further investigations using gradient-based approaches have revealed a progressive decoupling of principal functional and microstructural gradients,^12^ indicating the flexible functional roles of transmodal networks.^46^ Gradient techniques, therefore, unify different principles of brain organization across multiple neurobiological features and scales.

Neurosciences has increasingly benefitted from and embraced open science practices, particularly through data sharing initiatives and the dissemination of derivative data alongside the publication of processing pipelines. Large collaborative projects have produced open source datasets acquired at 7T, such as the Human Connectome Project.^22^ However, these datasets focused either on in-depth sampling of functional scans^22^ or structural datasets, to mainly explore subcortical structures.^21^ To fill this gap, we provide the Precision Neuroimaging and Connectomics (PNI) dataset, which capitalized on 7T MRI acquisitions across multiple sessions. This dataset offers significant advancement by providing a multimodal, multi-sequence, and multi-session MRI resource, specifically designed to enable ultra-high-resolution, multimodal precision mapping of the human brain at the 7T. This dataset includes anonymized raw data that conforms with Brain Imaging Data structure^47^ (BIDS) standards and processed data derivatives using an open access pipeline^48^, which include inter-regional connectomes derived from multi-state fMRI, diffusion tractography, multiple quantitative imaging for microstructure covariance analysis and geodesic cortical distances that are constructed across multiple spatial and parcellation schemes. By providing this multimodal precision neuroimaging dataset, we enable a level of human brain mapping that provides novel insights, that are not readily achievable with current open-source datasets. Our initiative promises to become an invaluable and openly accessible resource for researchers aiming to advance our understanding of structure-function relationships in the human brain.

## Methods

### Participants

The 7T MRI protocol was implemented at the McConnell Brain Imaging Centre of the Montreal Neurological Institute (The Neuro) between March 2022 and November 2023. Ten healthy adults (4males/6females, age=26.6±4.60, left/right-handed=2/8) with no signs of neurological or psychiatric illness underwent three testing sessions (interval between sessions: mean±SD=95.45(74.71 days)). The MRI data acquisition protocols were approved by the Research Ethics Board of McGill University and the Montreal Neurological Institute (2023-8971/2022- 8526). All participants provided written informed consent, which included a provision for openly sharing all data in anonymized form.

### MRI data acquisition

MRI data were acquired on a 7T Terra Siemens with a 32-receive and 8-transmit channel head coil in parallel transmission (pTX) mode. Participants underwent 4 distinct structural and 5 distinct functional imaging protocols across three different sessions, with total scanning time of ∼90 minutes/session (**Figure 1**).

**Figure.1.**
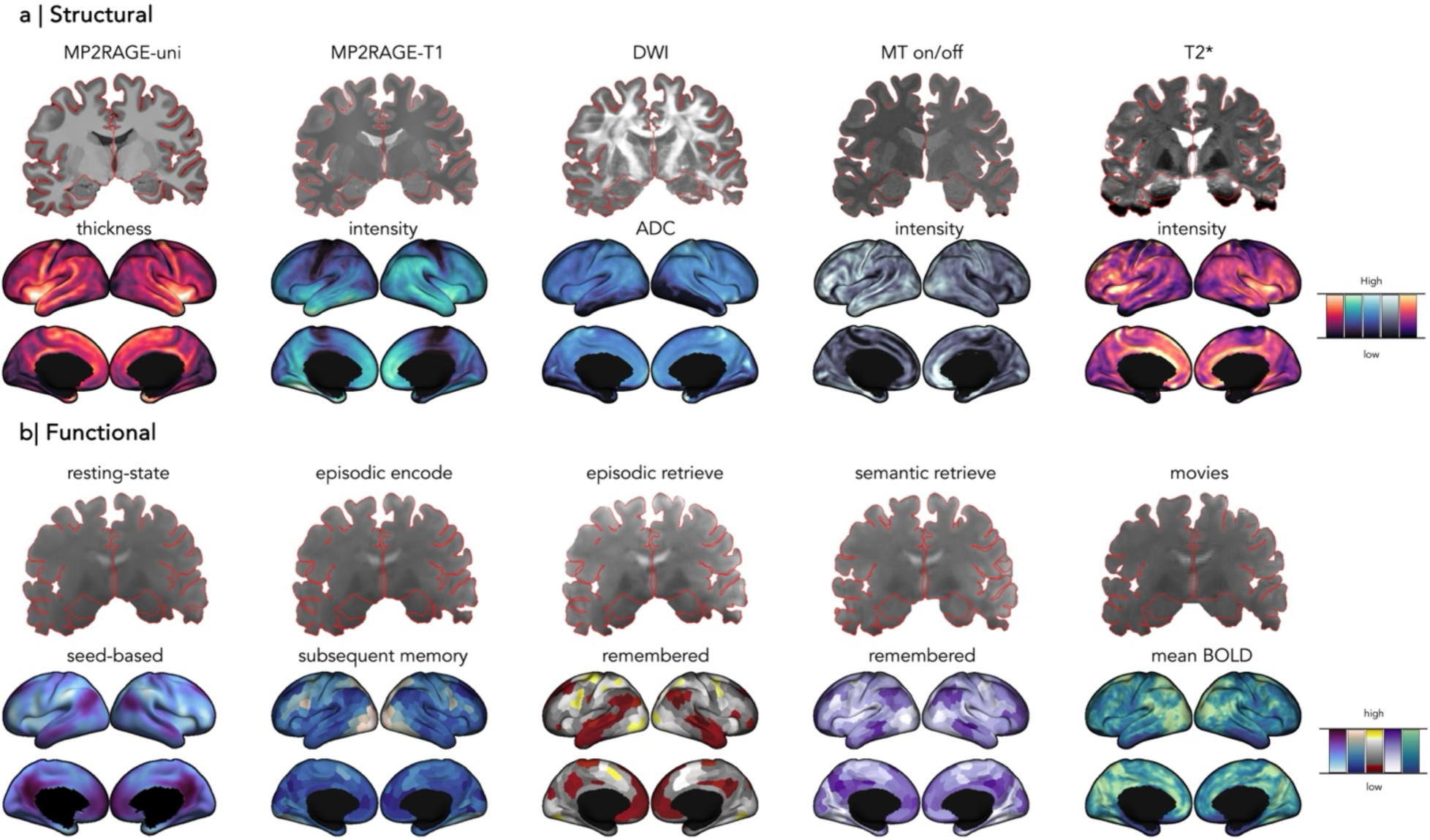
Precision NeuroImaging and Connectomics (PNI) dataset. Multimodal structural (**a)** and functional MRI (**b**) sequences acquired across multiple sessions and below are example neuroimaging features that can be derived from each modality. Cortical thickness and T1 map intensity features were generated from MP2RAGE-uni and T1 map, respectively. ADC was from the DWI. For MT on/off, the intensity feature was derived from magnetization transfer saturation (MTSAT). T2* map was first generated from fitting the T2*-weighted data, followed by intensity map. **Abbreviations:** MP2RAGE=3D-magnetization-prepared 2-rapid gradient-echo sequence, DWI=diffusion weighted imaging, MT=magnetization transfer, ADC=apparent diffusion coefficient, BOLD=blood oxygenation level dependent.

#### Structural imaging

Structural scans included: *(i)* three 3D-magnetization-prepared 2-rapid gradient-echo sequence with Universal Pulses to optimize B1^+^ uniformity^49^ (MP2RAGE; 0.5mm isovoxels, matrix=320×320, 320 sagittal slices, TR=5170ms, TE=2.44ms, TI1=1000ms, TI2=3200ms, flip=4°, iPAT=3, partial Fourier=6/8, FOV=260×260mm^2^), for studying cortical morphology and intracortical microstructural organization, *(ii)* two diffusion-weighted MRI scans with three distinct shells with b-values 0, 300, 700, and 2000s/mm^2^, with each shell acquired with 10, 40, and 90 diffusion weighting directions, respectively, (1.1mm isovoxels, TR=7383ms, TE=70.60ms, flip angle=90°, refocusing flip angle=180°, FOV=211×211mm^2^, slice thickness=1.1mm, MB=2, echo spacing=0.79ms) for examining structural connectomes and fiber architectures, *(iii)* one myelin-sensitive magnetization transfer (MT; 0.7mm isovoxels, TR=95ms, TE=3.8ms, flip angle=5°, FOV=230×230 mm^2^, slice thickness=0.72mm, 240 sagittal slices) and (*iv*) one iron-sensitive T2*- weighted multi-echo gradient-echo (ME-GRE) (0.7mm isovoxels, TR=43ms, TE=6.46-11.89- 17.33-22.76-28.19-33.62ms, flip angle=13°, FOV=240×240 mm^2^, slice thickness=0.72mm, 160 sagittal slices).

#### Functional imaging

All multi-echo fMRI were acquired with a 2D BOLD echo-planar imaging sequence^50^ (University of Minnesota, CMRR; 1.9mm isovoxels, 75 slices oriented to AC-PC-39 degrees, TR=1690ms, TE=10.80/27.3/43.8ms, flip=67°, FOV=224×224mm^2^, slice thickness=1.9mm, MB=3, echo spacing=0.53ms). During the *(v)* three resting-state fMRI sessions, participants were instructed to fixate on a grey cross and not think of anything for a duration of 6 minutes (210 time points). Task-based fMRI were implemented based on a validated open-source protocol,^17,51^ and included the *(vi/vii)* one session of episodic encoding/retrieval and *(viii)* semantic tasks, each lasting approximately six minutes. During the episodic memory encoding, participants memorized paired images of objects. In the retrieval phase, participants were shown an image and asked to identify the paired object from three options. Semantic memory retrieval involved identifying the object that is most conceptually related to a target image from three options. In both memory tasks, there were 48 trials and the difficulty was modulated based on semantic relatedness scores,^52^ ensuring a balanced difficulty level across trials (*i.e.,* with 24 difficult and 24 easy trials). We also collected fMRI data while participants watched affective and documentary-style movies *(ix)*, allowing to track hemodynamic activity during naturalistic viewing conditions.^18,53^ A detailed description of the imaging protocol is additionally provided in this data release, which includes the complete list of acquisition parameters.

### MRI data preprocessing

Raw DICOMS were sorted according to sequence and converted to Nifti format using dcm2niix^54^ (https://github.com/rordenlab/dcm2niix). Subject and session-specific directories were created according to BIDS convention^55^ (https://bids.neuroimaging.io) and validated with the BIDS validator^47^ v1.13 (https://doi.org/10.5281/zenodo.3762221). All functional and structural data were preprocessed with micapipe^48^ v.0.2.3 (http://micapipe.readthedocs.io), an open-access preprocessing software. Structural MRI images were anonymized and defaced^48^.

#### T1w processing

The MP2RAGE-derived uniform images (*uni*) were initially deobliqued and adjusted to LPI orientation (*left to right, posterior to anterior, and inferior to superior*). Subsequently, background denoising,^56^ bias-field correction and intensity normalization^57^ were applied. The skull-stripped image and subcortical structures were segmented using FSL FIRST.^58^ To improve image contrast, the image underwent additional non-local means filtering,^57,59^ followed by generation of cortical surface segmentations using FastSurfer^60^ v.2.0.0. Visual inspection and additional quality control quantifications and corrections ensured the accuracy of resulting surface outputs.

#### Quantitative image processing

Traditional quantitative maps require correction for bias introduced by RF transmit field (B1+) inhomogeneities,^61,62^ a process typically requiring fitting the B1+ mapping. To address this, we implemented a unified segmentation-based correction method^62^ (UNICORT; kernel=20mm full-width-at-half-maximum, FHWM, normalization parameter=’extremely light’) to all the raw images. UNICORT employs a probabilistic framework that incorporates a physically informed generative model of smooth B1+ inhomogeneities and their multiplicative effect on quantitative maps, resulting in improve data quality and efficiency. The T2* maps were generated by fitting the T2*-weighted data and were bias-corrected. T2* maps were additionally denoised using an adaptive optimized non-local means^63^ (AONLM) filter (patch size=3×3×3voxels, search size=7×7×7voxels; beta=1.0). This filter operates on patches within the image to perform denoising, effectively removing spatially varying noise introduced by the GRAPPA technique. MT saturation^61^ (MTSAT), a semi-quantitative metric that represents the proportion of free water saturated by a single MT pulse within repetition time, was generated from bias-corrected MT images using qMRLab^61^ (https://qmrlab.readthedocs.io). Finally, subject and quantitative-image specific series of equivolumetric surfaces between pial and white matter boundaries were constructed, resulting in a unique intracortical intensity profile at each vertex. Each quantitative image was then aligned to native FastSurfer space of each participant using label-based affine registration.^64^ No further processing was applied to the quantitative images.

#### Diffusion MRI processing

Session-specific diffusion MRI data were concatenated and preprocessed in native diffusion MRI space with MRtrix.^65^ The processing included Marchenko-Pasteur^66,67^ denoising, correction for Gibbs ringing artifact,^68,69^ head motion, susceptibility distortion, eddy current-induced distortion and motion, as well as non-uniformity bias field correction.^57,69–71^ Subsequently, the b0 image was extracted and linearly registered to the main structural image (*i.e.,* MP2RAGE-*uni*). Finally, fractional anisotropy and mean diffusivity maps,^72^ considered as surrogates of fiber architecture and tissue microstructures, were computed by fitting a diffusion tensor model.^73^

#### Multi-echo fMRI data processing

Resting-, task- and movie-state fMRI were processed using a combination of FSL^74^ 6.0, , AFNI^75^ 20.3.0 and ANTs^76^ 2.3.4 software. Initially, each echo was reoriented to LPI, and motion corrected. Multi-echo scans underwent further processing with TEDANA^77^ v.0.0.12 ( https://tedana.readthedocs.io/), integrated within the *micapipe*^48^ processing framework (https://micapipe.readthedocs.io/en/latest/pages/02.restingstateproc/index.html). The TEDANA pipeline extracts time series from all echos, optimally combines them, and decomposes the multi-echo BOLD data via principal and independent component analysis. TE-dependent components were classified as BOLD, and independent components discarded. Data then underwent high-pass filtering, followed by registration of volumetric time series to the native cortical surface. Additionally, native surface time series were registered to different surface templates (*i.e.,* fsLR- 32k, fsaverage5), followed by correction for motion spikes and global signal using linear regression. Cerebellar and subcortical timeseries were also included in this release.

### Vertex-wise individual and group-level connectome matrices

Inter-regional structural and functional connectomes derived from each imaging sequence are also included in this data release (**Figure 2**). All atlases, available on fsLR-32k symmetric surface template,^78^ were resampled to each subject’s native surface for modality-, subject-, and session-specific matrix generation. Neocortical connectomes from 18 distinct parcellations including: *(i)* anatomical atlases from Desikan-Killiany^79^ (*aparc*), Destrieux^80^ (*aparc.a2009s*), an *in-vivo* approximation of cytoarchitectonic parcellation studies by Von Economo and Koskinas,^81^ and sub-parcellations within Desikan-Killiany^79^ atlas (100-400 parcels); *(ii)* intrinsic functional connectivity based parcellations^82^ (Schaefer atlases based on 7-network parcellation) ranging from 100-1000 nodes; and *(iii)* multimodal parcellation atlas^83^ from the Human Connectome Project with 360 nodes (Glasser parcellation). Connectome matrices furthermore encompass data for hippocampus and subcortical structures including the nucleus accumbens, amygdala, caudate nucleus, pallidum, putamen, and thalamus.

**Figure 2.**
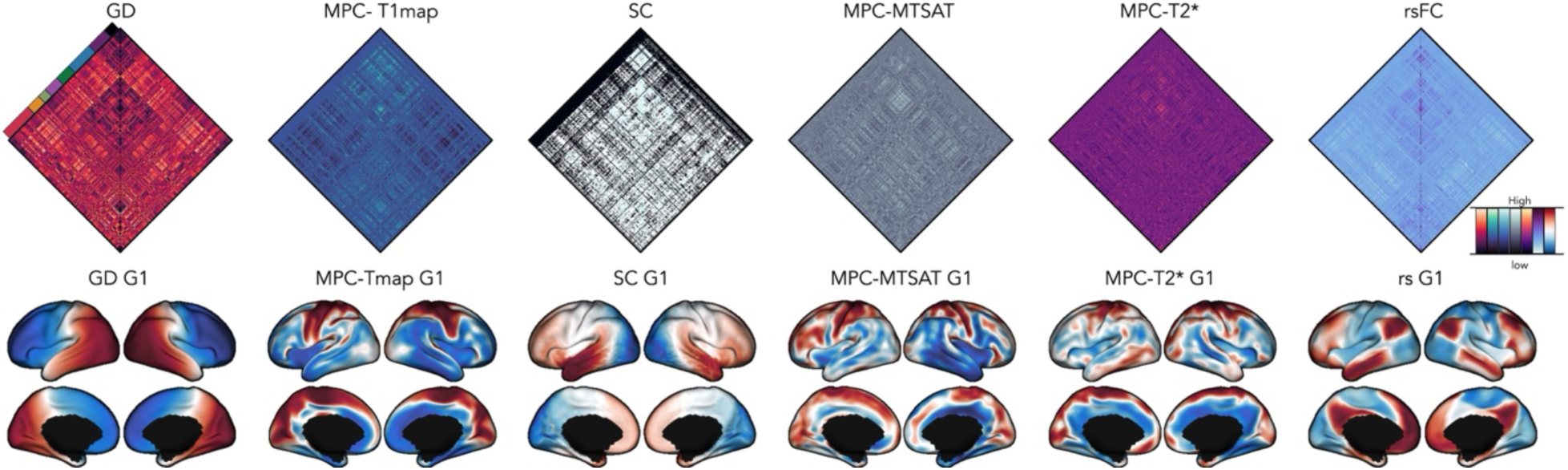
Connectomes and Gradients. Vertex-wise connectomes sorted according to the Yeo-7-network parcellations^90^ and principal gradients derived from different modalities are included in this release. **Abbreviations:** GD=geodesic distance, MPC=microstructural profile covariance, SC=structural connectome, MTSAT=magnetization transfer saturation, rs=resting state, FC=functional connectome, G1=Gradient 1.

Geodesic distances (GD) between all cortical parcels on the subject’s native midsurface were computed using Dijkstra’s algorithm.^84^ The central vertex for each parcel, identified as the vertex with the shortest summed Euclidean distance to all other vertices within the parcel, was used as a reference point. The GD from the centroid vertex to all other vertices on the midsurface mesh was computed using workbench.^85^ Distances were averaged within each parcel.

Microstructure profile covariance (MPC) matrices were individually computed from each quantitative MRI contrast (*e.g.,* MP2RAGE-derived T1map, MTSAT, T2*map). Initially, intracortical intensities were sampled by constructing 16 equivolumetric surfaces between pial and white matter boundaries.^86^ Boundary surfaces were discarded to account for partial volume effect. The resulting 14 profiles were used to compute MPC matrices, which captures the similarity in intracortical microstructure across cortical regions. Specifically, vertex-wise intensity profiles are averaged within parcels for each parcellation, and nodal profiles are cross-correlated across the cortex using partial correlation, while controlling for the average cortex-wide intensity profile. Notably, regions such as the left/right medial walls, corpus callosum, and pericallosal regions are excluded when averaging cortex-wide intensity profiles.

Structural connectomes (SC) were generated using Mrtrix3 from preprocessed diffusion MRI data, with subcortical and cerebellar parcellations registered to native diffusion MRI space. Initially, a tractogram was generated using the iFOD2 algorithm and 3-tissue anatomically constrained tractography^87,88^ (cortical and subcortical grey matter, white matter, cerebrospinal fluid), producing 40M streamlines (maximum tract length=400, minimum length=10, cutoff=0.06, step=0.5). Tractograms underwent spherical deconvolution informed filtering^89^ to reconstruct whole brain streamlines that are weighted by cross-sectional multipliers on diffusion MRI native space. Connection strengths between nodes were calculated based on the weighted streamline count and edge length matrices were subsequently generated.

Functional connectivity (FC) matrices were individually computed from each fMRI scan. Session- and state-specific time series were mapped to individual surface models and registered to standard templates (*e.g.,* fsLR-32k). Within cortical parcels, the native surface and fsLR-32k surface-mapped timeseries were averaged. Subcortical and cerebellar parcellations were warped to each participant’s native fMRI volume space for nodal time series extraction. Finally, subject- and state-specific functional connectomes were generated by cross-correlating all nodal time series.

## Data Records

All files are organized and conform with BIDS^55^ and are available on the Open Science Framework^91^(OSF; https://osf.io/mhq3f/). The raw data for each participant totals ∼14 GB, while subject-specific derivatives are ∼ 46 GB. Due to storage limitations on the OSF platform, raw data and derivatives were uploaded as separate project components. The raw data and processed files were further compressed into at least 3-subject and 12-file batches, respectively (see README files for complete documentation).

### Raw data

All data in native space and .*json* sidecars are located in */rawdata/sub-PNC#/ses-#* branch of the BIDS directory structure. Each subject underwent three-sessions of MP2RAGE sequence imaging and rs-fMRI (/*ses-01, /ses-02, /ses-03*), two-sessions of DWI (*/ses-01, /ses-02*), single session of multitask-based (/*ses-01 or /ses-02*) and movie-fMRI, as well as MT- and T2*-weighted imaging (/*ses-03*; **Figure 4**).

For each subject and session (*/sub-PNC#/ses-#*), all defaced structural files are located in the */anat* directory: MP2RAGE-derived T1w images (denoted as *uni*), inversion time parameters (*inv-1, inv-2*) and T1 relaxometry (*T1map*). The DWI files are contained in /*rawdata/sub-PNC#/ses-#/dwi* subdirectory which included: diffusion gradient and direction (*bval, bvec*), DWI volumes and .json files associated with each shell (*sub-PNC#_ses-#_acq-b#_dir-AP_dwi.json*).

The */sub-PNC#/ses-#/fmap* subdirectory contains b0 images in inverse phase encoding direction (*i.e., sub-PNC#_ses-0#_acq-fmri_dir-AP_epi.nii.gz*) and phase encoding direction of spin-echo images (*i.e., sub-PNC#_ses-#_acq-fmri_dir-PA_epi.json, sub-PNC#_ses-#_acq-fmri_dir-PA_epi.json*)

Subject and session-specific multi-echo fMRI scans and corresponding *tsv* event files for tasks are located in /*rawdata/sub-PNC#/ses-#/func* subdirectory. Functional timeseries for all rs-fMRI include 210 time points (*i.e., sub-PNC#_ses-#_task-rest_echo-#_bold.nii.gz*), episodic encoding task with 200 timepoints (*i.e., sub-PNC#_ses-#_task-epiencode_echo-#_bold.nii.gz*), episodic retrieval with 205 timepoints (*i.e., sub-PNC#_ses-#_task-epiretrieve_echo-#_bold.nii.gz*), semantic retrieval blocks 1 and 2 (*i.e., sub-PNC#_ses-#_task-semantic#_echo-#_bold.nii.gz*), each with 125 timepoints, two affective and two documentary style movies, each with 105 time points (*i.e., sub-PNC#_ses-#_task-movies#_echo-#_bold.nii.gz*).

### Processed data

Processed data are located in the */derivatives* subdirectory. Quality control reports for raw structural and the optimally combined echo of functional data are provided in the */derivatives/mriqc*/ directory. Image modality-specific matrices with node counts ranging from 70 to 1000 were generated using micapipe^48^, and are stored within their respective subdirectories (*e.g.,* structural connectomes can be found in */derivatives/micapipe_v0.2.0/sub-PNC#/ses-#/dwi/connectomes*, whereas functional connectomes can be found at */derivatives/ micapipe_v0.2.0/sub-PNC#/ses-#/func/desc-me_task-#_bold connectomes/surf*).

#### Structural data

Processing derivatives of structural scans are provided in /*derivatives/micapipe_v0.2.0/sub-PNC#/ses-#/anat*, which include the main structural scan (*i.e., sub-PNC#_ses-#_space-nativepro_T1w_nlm.nii.gz, nlm* is added to the string name to denote the non-local means filtering applied to the data). Furthermore, /*derivatives/micapipe_v0.2.0/sub-PNC#/ses-#/dist* contains the GD matrices for each cortical parcellation/surface (*i.e., sub-PNC#_ses-#_atlas-schaefer-1000_GD.shape.gii*) which was computed along each participants’s native midsurface using workbench command. Finally, the MPC matrices and intensity profiles generated from each quantitative image are stored in */derivatives/micapipe_v0.2.0/sub-PNC#/ses-#/mpc* (*i.e., acq-T1map, acq-MTSAT*) and identified by parcellation scheme from which they were computed (*i.e., sub-PNC#_ses-#_atlas-schaefer -400_desc-intensity_profiles.shape.gii, sub-PNC#_ses-#_atlas-schaefer -400_desc-MPC.shape.gii*).

#### Diffusion MRI data

Processing derivatives of diffusion MRI scans are provided in /*derivatives/micapipe_v0.2.0/sub-PNC#/ses-#/dwi* and organized into two distinct subdirectories. First, the structural connectomes and associated edge lengths are provided for each parcellation (*i.e., /dwi/connectomes/sub-PNC#_ses-#_space_dwi_atlas-schaefer-600_desc-iFOD2-40M-SIFT2_full-connectome.shape.gii*). Eddy outputs estimated and used for correcting eddy currents and movements are in */derivatives/micapipe_v0.2.0/sub-PNC#/ses-#/dwi/eddy*.

#### Multi-echo fMRI data

Functional MRI-specific (i*.e., desc-me_task-rest_bold, desc-me_task-epiretrieve_bold*) are provided in */derivatives/micapipe_v0.2.0/sub-PNC#/ses-#/func.* Each fMRI subdirectory is organized into two distinct subdirectories: *(1) /func/surf* which include all the fully processed connectomes that are computed from native-surface mapped timeseries as well as the temporal signal to noise ratio (*tSNR*) maps. *(2) /func/volumetric/* which mainly include the fully processed data in volumetric space (*i.e., sub-PNC#_ses-#_space-func_desc-me_preproc.nii.gz*).

#### Quality Control

Image quality matrices (IQM) computed by MRIQC v23.1.0 (https://github.com/nipreps/mriqc) are also provided in */mriqc* branch of PNI processing derivatives. Individual and session-specific IQM reports for structural (*/mriqc/sub-PNC#/ses-#/anat*), DWI (*/mriqc/sub-PNC#/ses-#/dwi*) and functional scans (*/mriqc/sub-PNC#/ses-#/func*) are provided in both *.html* and *.json* formats. These reports evaluates the quality of the input data such as contrast-to-noise ratio estimates and motion.^92^ The group-level quality control for the *micapipe* outputs *(/derivatives/micapipe_v0.2.0/micapipe_group-QC.pdf*) includes module processing tables and progress graphs and subject-wise processing times. It also covers key metrics for structural, diffusion, functional and quantitative MRI data such as mean cortical thickness and similarity matrices.

### Technical Validation

#### Cortical surface segmentations

All surface extractions were visually inspected by two raters (DGC, YW) and corrected for any segmentation errors with manual correction.

#### MRI image quality metrics

The MP2RAGE-uni, MT on/off and T2*-weighted image quality were evaluated using contrast-to-noise (CNR)^93^ derived with MRIQC.^92^ This metric relates to the separability between grey and white matter distributions in each image across all sessions (**Figure 3a**). For diffusion MRI images, total movement in each volume was quantified in each shell using MRtrix and FSLeddy,^70^ by calculating the displacement of each voxel and then averaging the squares (RMS) of those displacements across all intracerebral voxels. Additionally, diffusion MRI image quality was evaluated with MRIQC^92^ derived metrics, including SNR within the corpus callosum, framewise displacement (FD) and the presence of global spikes (**Figure 3b**). For each of the five functional scans, FD for the optimally combined echo was estimated using the FSL motion outlier detection tool. The tSNR for each subject and functional scan was computed by dividing the mean timeseries by the standard deviation. This was performed on the minimally processed timeseries to generate tSNR maps across the cortex for each subject. These native cortical surface timeseries were then coregistered to fsLR-32k surface templates and averaged across subjects and sessions (**Figure 3c**).

**Figure 3.**
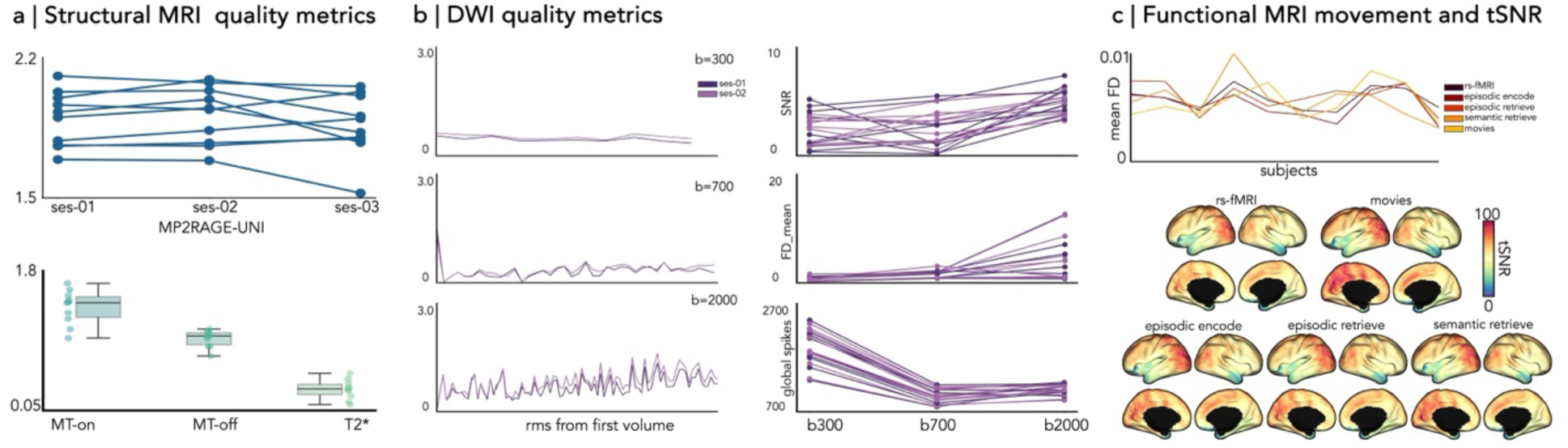
Technical validation metrics. (**a**) Image quality was evaluated with contrast-to-noise (CNR) estimated with the MRIQC pipeline (plotted for each individual across three sessions). No outliers were detected in MP2RAGE- uni scans across all sessions (upper panel), MT on/off (middle), and T2* (lower). (**b, left**) Motion parameters of diffusion-weighted images obtained from FSL eddy. Line plots illustrate root mean squared (RMS) voxel-wise displacement relative to the first volume across all shells, averaged across all subjects. (**b, right**) Individual signal-to-noise ratio (SNR) in the corpus callosum, framewise displacement (FD) and global spikes across all shells are also plotted. (**c, upper panel**) Framewise displacement (FD) of optimally combined echo for each functional scans obtained using FSL motion outliers, representing the average rotation and translation parameter differences at each volume. (**c, lower**). Vertex-wise temporal signal-to-noise (tSNR) computed on the native surface of each participant. Computed tSNR values were averaged within a fsLR-5k-node functional atlas and across individuals.

**Figure 4.**
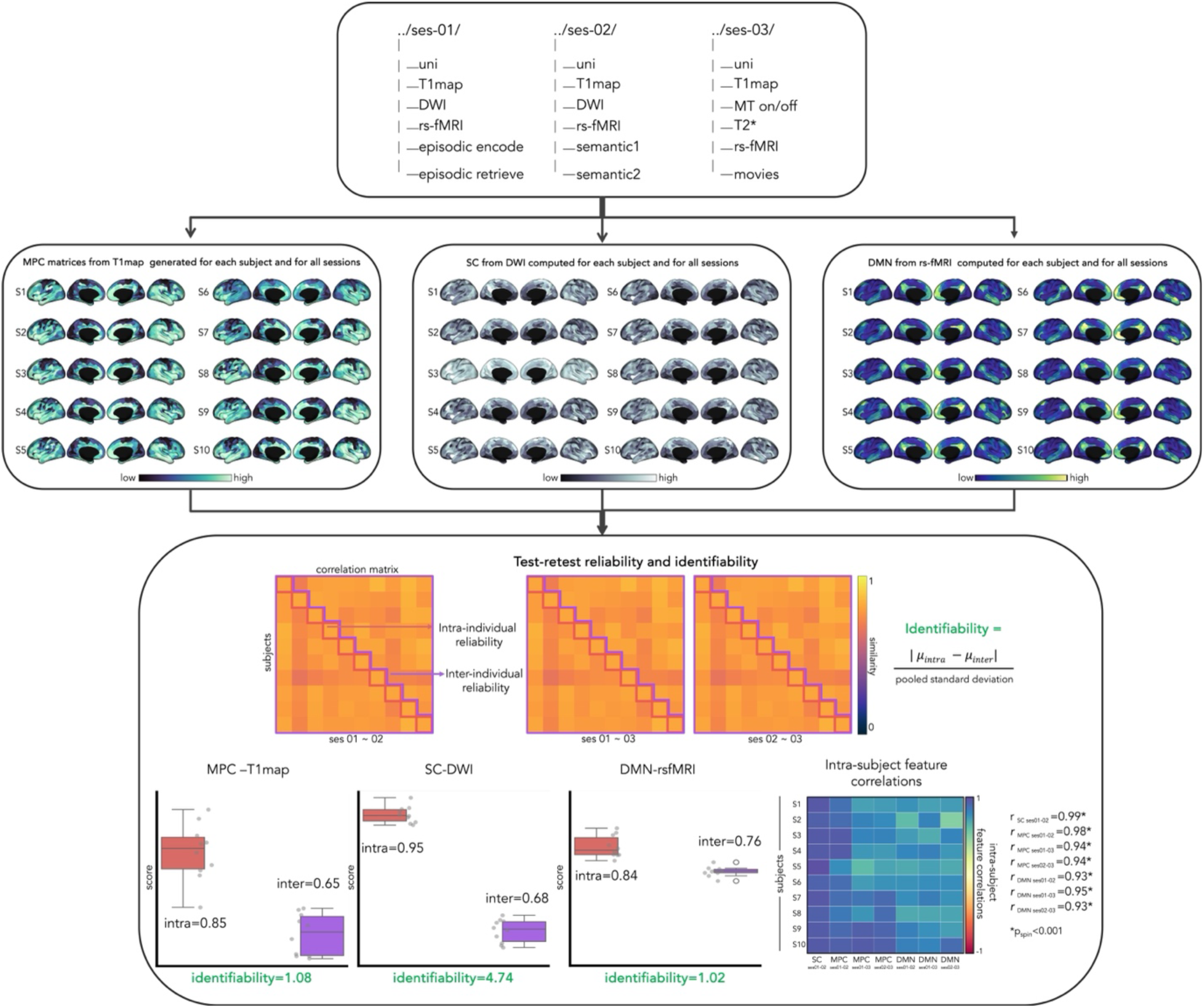
Identifiability and Reliability. Multimodal structural and functional MRI acquired across three sessions (**top panel**). Subject-specific MPC profiles derived from MP2RAGE-T1 maps, SC from DWI and DMN derived from resting state connectome (**middle panel**). Test-retest reliability and identifiability for MPC-T1maps, SC and DMN and within-subject feature correlations for each session (**bottom panel**). **Abbreviations:** DWI=diffusion weighted imaging, MT=magnetization transfer on/off, rs-fMRI=resting state functional MRI, S_j_=subjects, MPC=microstructural profile covariance, SC=structural connectome, DMN= Default mode network.

#### Surface-based neuroimaging features

Preprocessed data for each modality includes various surface-based neuroimaging features (**Figure 1**). Morphological features such as cortical thickness and curvature, as well as diffusion MRI derived FA and ADC are in */derivatives/micapipe_v0.2.0/sub-PNC#/ses-#/maps.* Tissue contrast maps are also available *(e.g., /derivatives/micapipe_v0.2.0/sub-PNC#/ses-#/mpc/acq-T1map/sub-PNC#_ses-01_atlas-schaefer-400_desc-intensity_profiles.shape.gii).* Functional connectivity matrices are available for each of the functional scans. All features are available in both native and standard surface templates (*e.g.,* fsLR-32k, fsaverage5). Additionally, we performed surface-based first level analysis to illustrate general linear model (GLM) fitting to task-based data sampled on the cortical surface.

#### Cortical gradient estimation from vertex-wise connectomes

Individual and group-level connectivity gradients were derived from each data modality using Brainspace v0.1.3 (https://brainspace.readthedocs.io/).^33^ Group-level gradients were generated using averaged subject-level matrices in fsLR-32k surface space (**Figure 2**). GD, MPCs and FC matrices were thresholded to retain only the top 10% row-wise connections. To reduce variance in connectivity strength, SC matrices were initially log-transformed before averaging. To mitigate limitations in mapping inter-hemispheric fibers with diffusion tractography, SC gradients were computed separately for each hemisphere and subsequently aligned the right to the left hemispheres. GD and corresponding gradients were also computed separately for each hemisphere. We constructed affinity matrices using the normalized angle to measure the similarity of inter-regional patterns between regions. Affinity matrices from each modality were fed into diffusion map embedding,^31,33,94^ a non-linear dimensionality reduction technique to identify low-dimensional eigenvectors. To evaluate reproducibility, individual and modality-specific gradients were also generated. Individual gradients were aligned to a group-level template using Procrustes rotations to ensure consistency in ordering and polarity across participants and modalities. This alignment was essential, as individual gradients may vary in orientation and magnitude, and aligning them to a common template enables meaningful group-level comparisons.^95^ Finally, we computed the averaged correlations between individual and group level gradients. The three group and individual-level gradients that explained most of the variance are provided in */derivatives/gradients*.

As expected, the principal GD gradient (GD-G1) recapitulated the longest cortical distance axis in anterior to posterior direction^1^. SC-G1 distinguished visual and sensorimotor surfaces. Notably, SC-G1 from Park and colleagues^34^ revealed a different pattern, distinguishing sensorimotor regions from the medial prefrontal anchor, but still overlaps with our findings in which the same pattern could be observe in our SC-G3 (see */derivatives/gradients*). The first three structural gradients, explaining a total of 45% of variance, captures within hemispheric visual temporo-occipital (SC-G1), fronto-parietal (SC-G2) and sensorimotor regions (SC-G3). Resting-state FC- G1 describe a unimodal to transmodal pattern.^31^ MPC-G1 derived from T1, MTSAT and T2* maps were anchored in primary sensory areas and limbic regions. ^12,14,41^ G1-GD (multi-session mean *r* and SD=0.99 ± 0.001; single session=0.99 ± 0.001), MPC-T1 (0.85±0.022; 0.79±0.04), and FC (0.86±0.04; 0.76±0.06) were highly replicable in all participants and moderately replicable for G1 SC (0.73±0.068; 0.66±0.13) and MPC-MTSAT (0.56±0.140) and MPC-T2* (0.40±0.423).

### Identifiability and Reliability

Our multi-session data allows to assess the test-retest reliability (**Figure 4, top**). For illustration, we examined the default mode network (DMN) connectivity from rs-fMRI connectomes, structural connectomes (SC) from diffusion MRI images and MPC profiles from MP2RAGE- derived T1 maps for individual and across three-scanning sessions for each of the 10 participants (**Figure 4 middle**). To assess the consistency of different modalities, we computed intra-subject feature correlations across sessions, revealing high consistency (*p_spin_*<0.001). We evaluated the reliability using a statistical framework^96^ that considers both intra- and inter-subject reliability. Intra- and inter-subject reliability were assessed by averaging the correlations between measurements obtained at each session and across participants, respectively. Ideally, individual MRI features should exhibit high reliability indicating consistency and lower population reliability preserving individual differences. Additionally, we evaluated the uniqueness of individual features using an established identifiability framework,^96–98^ measuring the effect size of differences between intra- and inter-individual reliabilities. Our analysis showed that for all features, the intra-subject (DMN=0.84, SC=0.95, MPC=0.85) was higher than inter-subject reliability (DMN=0.76, SC=0.68, MPC=0.65), with strong identifiability (DMN=1.02, SC= 4.74, MPC=1.08), indicating reliable and distinct DMN, SC and MPC patterns from our UHF data while preserving individual differences (**Figure 4 bottom**).

## Code Availability

The processing pipeline scripts, including usage instructions and processing steps are openly available in GitHub (https://github.com/MICA-MNI/micapipe) and *ReadTheDocs* (https://micapipe.readthedocs.io/). Gradients were generated using Brainspace v0.1.3 (https://brainspace.readthedocs.io/).

## Acknowledgements

The authors thank all participants who took part in this multi-session study. We also sincerely thank David Costa, Ronald Lopez, Soheil Quchani, and Michael Ferreira for their assistance in data collection. DGC received support from the Fonds de la Recherche du Québec – Santé (FRQ- S), Quebec BioImaging Network (QBIN), Savoy Foundation, Brain Canada, and Canada First Research Excellence Fund, awarded to McGill University for the Healthy Brains for Healthy Lives (HBHL) initiative. JR received support from the Canadian Open Neuroscience Platform (CONP) and Canadian Institute of Health Research (CIHR). CP and RRC received support from the FRQ-S. JD is supported by Natural Sciences and Engineering Research Council-Post-Doctoral Fellowship (NSERC-PDF). NE is supported by Sir Henry Wellcome Postdoctoral Fellowship from the Wellcome Trust [222799/Z/21/Z]. ST received a Faculty of Medicine studentship from McGill University. OB received support from the HBHL program. YW received support from the FRQ- Nature et technologies (FRQ-NT). BCB acknowledges support from CIHR (FDN-154298, PJT- 174995, PJT-191853), SickKids Foundation (NI17-039), NSERC (Discovery-1304413) BrainCanada, the Helmholtz International BigBrain Analytics and Learning Laboratory (HIBALL), HBHL, the Canada Research Chairs Program, and the Centre for Excellence in Epilepsy at the Neuro (CEEN).

## Author contributions

Conception, design, manuscript preparation: DGC, BCB, IRL, JS, RRC; Participant recruitment: DGC; Data acquisition: DGC, IRL, ST, Processing pipeline: RRC, YH; Data processing: DGC, VK; Quality control: DGC, YW. All authors provided feedback and approved the final manuscript.

## Competing interests

The authors declare no competing interests.

## Notes

### Competing Interest Statement

The authors have declared no competing interest.

### Summary of Updates

Revised title and abstract to include the word MRI. Expanded the main goal of the data release.Restructured the Methods section for better flow. Included more quality control for the MRI images. Updated Figure 3 in the technical validation section to include DWI image quality control. Added justification on gradient differences from previous paper in the cortical gradient estimation from vertex wise connectomes section. Finally, the identifiability and realibility section was updated to include the analysis for structural connectomes.

